# Axon termination of the SAB motor neurons in *C. elegans* depends on pre- and postsynaptic activity

**DOI:** 10.1101/2025.09.05.674448

**Authors:** Alexis Weinreb, Minh-Dao Nguyen, Quentin Lemaitre, Laure Granger, Maelle Jospin, Jean-Louis Bessereau

## Abstract

Axon termination is a critical step in neural circuit formation, but the contribution of activity from postsynaptic targets to this process remains unclear. Using *Caenorhabditis elegans* SAB neurons as a model system, we showed that inhibition of muscle activity during a critical period of postembryonic development led to axonal overgrowth and ectopic synapse formation. This effect is mediated by a local retrograde signal and requires neuronal voltage-gated calcium channels (VGCCs) acting cell-autonomously to constrain axon growth. Manipulating SAB neuron excitability demonstrated that increased intrinsic neuronal activity drives overgrowth, while reducing activity suppresses it, establishing a functional link between muscle-derived cues and presynaptic excitability. Transcriptomic analysis and genetic studies further implicate the neuropeptides FLP-18 and NLP-12 as essential modulators of this activity-dependent process. Our findings reveal a temporally and spatially restricted retrograde signaling mechanism in motor neurons, where target activity, neuronal calcium dynamics and neuropeptide signaling cooperate to ensure proper axon termination. These results highlight conserved principles of activity-dependent regulation at neuromuscular junctions and provide a framework for understanding how motor circuits integrate target feedback to sculpt precise connectivity.

## Introduction

Axonal morphology is largely established during development, as axons extend, navigate toward their targets, and terminate at precise locations to form functional circuits. While the signaling mechanisms underlying axon growth and axon guidance have been extensively studied, the process of axon termination remains comparatively less understood. Beyond its fundamental biological interest, deciphering the molecular basis of axon termination has important implications for human health, since molecules involved in this process have been linked to neurodevelopmental disorders (AlAbdi et al., 2023; Buddell et al., 2019).

The nematode *Caenorhabditis elegans* is a powerful model system for investigating axon morphology due to its transparent body, stereotyped nervous system and versatile genetic toolkit (Nance and Frøkjær-Jensen, 2019). Studies in the touch receptor neurons ALM and PLM have uncovered multiple regulators of axon termination (Desbois and Grill, 2024), notably *rpm-1* and *fsn-1* (Borgen et al., 2017; Grill et al., 2007; Schaefer et al., 2000). *rpm-1* encodes a ubiquitin ligase of the well-conserved PHR family (Highwire in Drosophila, Phr-1 in mouse and Pam/MYCB2 in human) and forms a multi-subunit complex with the F-box protein FSN-1 (Liao et al., 2004). Interestingly, recent studies have revealed that the PLM and ALM axon termination defect observed in *fsn-1* can be suppressed by mutations in genes encoding voltage-gated calcium channel (VGCC) subunits (Buddell et al., 2019; Buddell and Quinn, 2021). These findings suggest that VGCCs negatively regulate an unidentified protein that cooperates with RPM-1 to promote axon termination. The effect of VGCCs could be mediated by modification of calcium fluxes or cell excitability.

Activity-dependent regulation of axon termination has been reported in a class of motor neurons in *C. elegans* called the SABs (Zhao and Nonet, 2000). The SABs form a set of three cholinergic motor neurons, whose cell bodies are located ventrally in the retrovesicular ganglion in the head of the worm (Figure 1A). SABVL and SABVR (**SAB v**entral **l**eft and **v**entral **r**ight) extend axons anteriorly, while SABD (**SAB d**orsal) initially projects dorsally before bifurcating into two anterior neurites. The four axons run along the head muscle cells toward the tip of the nose, and form *en passant* neuromuscular synapses. In their pioneering study, Zhao and Nonet (Zhao and Nonet, 2000) reported the following evidence: (i) in mutants with impaired synaptic transmission, SAB axons fail to terminate properly and display overgrowth phenotypes, such as sprouting or looping at the nose tip, (ii) reduction of muscle activity through ectopic expression of a gain-of-function potassium channel also induces axon termination defects and (iii) activity-dependent termination defects arise only when synaptic activity is disrupted during early larval stages.

**Figure 1.**
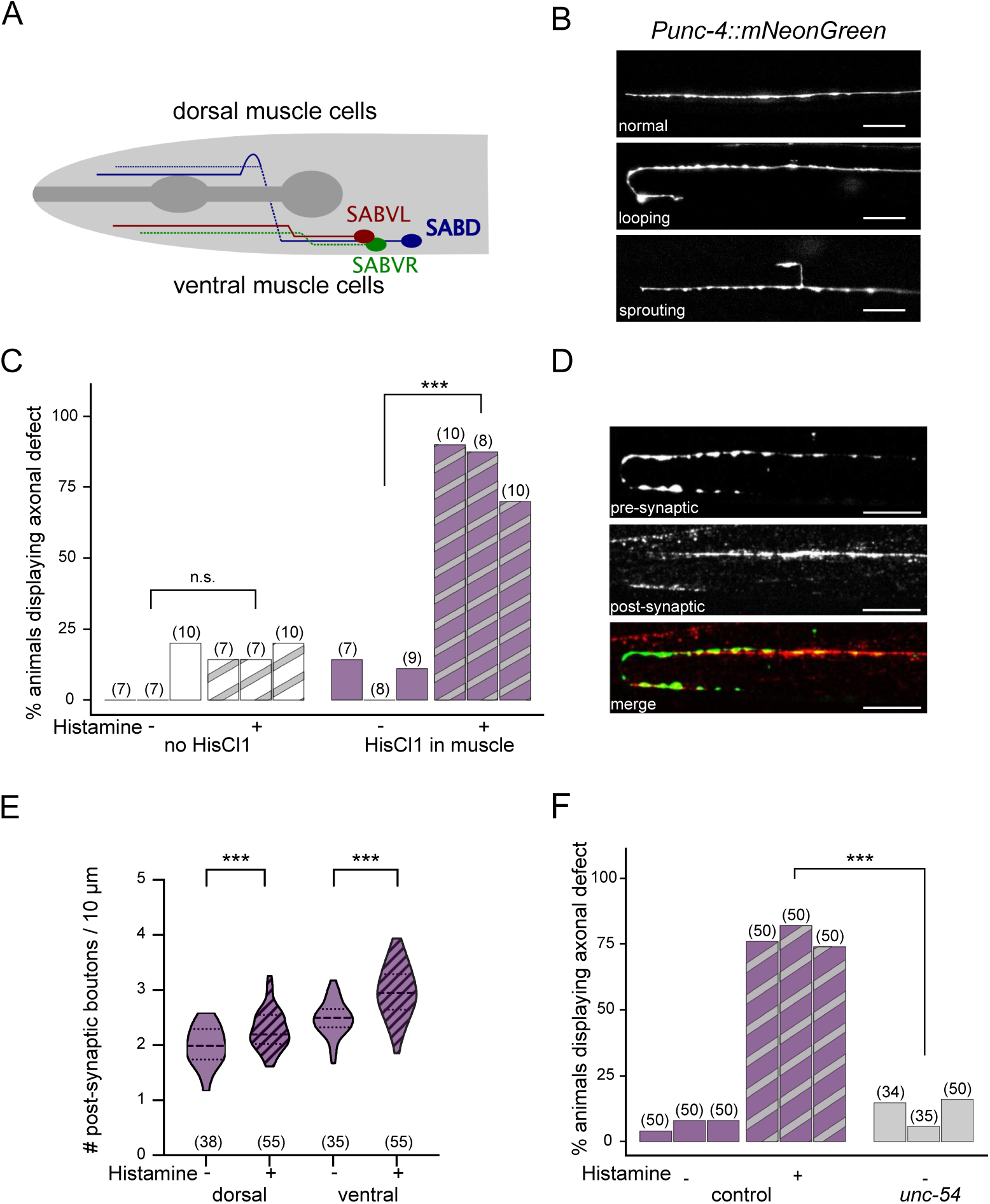
Reduced muscle electrical activity induces axonal overgrowth in SAB motor neurons. (**A**) Schematic of the three SAB motor neurons. The two ventral (SABV, left in red and right in green) and the dorsal neuron (SABD, in blue) have their cell bodies located in the retrovesicular ganglion, on the ventral side of the neck. Their axons project along head muscles (not represented), forming *en passant* synapses. (**B**) Confocal pictures of the head of animals carrying two single-copy integrated transgenes: *Punc-4::mNeonGreen* (to visualize SABs) and *Pmyo-3::HisCl1::SL2::tagBFP* (to silence muscles upon 10 mM histamine). Axonal morphology was classified as normal (top) or defective, either by exhibiting looping (middle) or sprouting (bottom). Nose is left, pharyngeal bulbs right. Scale bar: 10 µm. (**C**) Percentage of animals with SAB defects observed in animals either expressing *Punc-4::GFP* alone (no HisCl1) or together with *Pmyo-3::HisCl1* (HisCl1 in muscle, in purple), after water (-) or histamine (+) treatment (+). Cochran–Mantel–Haenszel (CMH) statistical tests. (**D**) Representative confocal pictures of presynaptic (*Punc-4::snb-1::GFP*, top) and postsynaptic (*unc-29(kr208::TagRFP-T*, middle) regions observed on a SAB axon exhibiting overgrowth defect. Merged image at bottom. Nose is left, pharyngeal bulbs right. Scale bar: 10 µm. (**E**) Violin plot of the density of postsynaptic boutons in animals expressing *Punc-4::mNeonGreen* and *Pmyo-3::HisCl1::SL2::tagBFP*, after water (-) or histamine (+) treatment. Dorsal and ventral axons were analyzed separately, numbers under violin plot indicate the number of axons scored coming from 18 animals treated by water and 27 treated by histamine. Mann-Whitney tests. Dashed lines, median; dotted lines, quartiles. (**F**) Percentage of animals with SAB overgrowth in control (*Pmyo-3::HisCl1::SL2::tagBFP*; *Punc-4::GFP*) or in *unc-54* mutant background, after water (-) or histamine (-) treatment. The CMH test compares the percentage of defects in control animals treated by histamine to *unc-54* mutants. In C and F, bars are grouped by experimental replicate (left to right). Numbers above bars indicate the number of animals scored. In all panels, n.s.: not significant, ***: p < 0.001.

To revisit and extend Zhao and Nonet’s findings on SAB axon termination, we generated single-copy transgenes that enable precise visualization of SAB axon morphology as well as spatial and temporal control of cell activity. We confirmed that reducing muscle activity during a critical developmental window induces SAB axon overgrowth, leading to the formation of ectopic synapses. We showed that local inhibition of head muscle activity alone was sufficient to impact SAB morphological development. Furthermore, we identified neuronal VGCCs as key mediators of this muscle activity-dependent remodeling and we provided evidence that SAB neuronal activity is required. Finally, we uncovered a role for neuropeptidergic signaling and pinpointed the neuropeptides *flp-18* and *nlp-12* as critical regulators of SAB axon overgrowth in response to decreased muscle activity.

## Results

### Muscle silencing induces axonal overgrowth in SAB motor neuron

We first aimed to reproduce the previously reported effect of muscle activity on SAB axonal morphology (Zhao and Nonet, 2000). To visualize the axons, we expressed the fluorescent protein mNeonGreen in SAB motor neurons under the control of a truncated version of *unc-4* promoter (see Methods). To inhibit muscle activity with spatial and temporal control, we generated a strain expressing a single-copy integrated transgene encoding the histamine-gated chloride channel (HisCl1) specifically in muscle cells (Pokala et al., 2014). Upon exposure to histamine, HisCl1 is activated, leading to chloride ion influx and membrane hyperpolarization in muscle cells. This treatment resulted in near-complete paralysis of transgenic worms, while wild-type animals showed no detectable effect (Supplementary Movies 1, 2).

In the absence of histamine, fewer than 10% of animals expressing HisCl1 exhibited axonal sprouting or looping, a rate similar to that of non-transgenic animals (Figure 1B, C). When HisCl1 was activated in muscle by histamine, quantification revealed a clear axonal overgrowth phenotype in SAB: more than 80% of the animals displayed axon sprouting or looping. Importantly, histamine had no effect in animals lacking HisCl1, validating the specificity of our assay (Figure 1C). Hereafter, we refer to animals expressing both mNeonGreen in SAB neurons and HisCl1 in muscle as the control genotype.

Varicosities were visible along the overgrown SAB axons when muscle activity was inhibited. To investigate whether these structures were associated with synapse formation, we built a strain expressing HisCl1 and tag-RFP-T-tagged acetylcholine receptors in muscle, along with GFP-tagged synaptobrevin-1 in SAB motor neurons. When the ectopic axonal branches grew in contact with muscle membrane, we frequently observed the accumulation of postsynaptic receptors near the ectopic varicosities (Figure 1D). We quantified the number of varicosities in control animals with or without histamine exposure and we found a significant increase in both dorsal and ventral SAB axons following histamine treatment (Figure 1E). This increase was consistent with formation of additional synapses along the overgrown axons.

We then asked whether SAB overgrowth was triggered by reduced muscle activity itself, or indirectly through complete paralysis and lack of movement. To address this, we examined SAB axons in *unc-54* mutants, which are paralyzed due to defects in Myosin Heavy Chain but retain normal level of muscle activity (Dibb et al., 1985). In these mutants, we did not observe an increase in SAB axonal overgrowth (Figure 1F). This result indicates that reduced muscle electrical activity, and not the absence of contractile activity, is the critical factor driving SAB axonal overgrowth.

### Overgrowth is induced by a local retrograde signal

SAB motor neurons only innervate the anterior-most muscle cells located in the head of the worm (Zhao and Nonet, 2000). In our previous experiments, muscle activity was inhibited throughout the entire body. To test whether inhibiting head muscle activity alone was sufficient, we drove the expression of HisCl1 specifically in head muscles using a fragment of the *tni-3/*troponin I promoter (Takashima et al., 2012). HisCl1 was strongly expressed in head muscles, with weaker expression in the vulva and stomato-intestinal muscles (Figure 2A). Histamine treatment efficiently and selectively silenced head muscles, evidenced by paralysis confined to the head and neck (Supplementary Movie 3). Under these conditions, 50% of treated animals exhibited SAB axonal overgrowth (Figure 2B). These results demonstrate that the signaling mechanism driving overgrowth involves, at least in part, a local retrograde signal.

**Figure 2.**
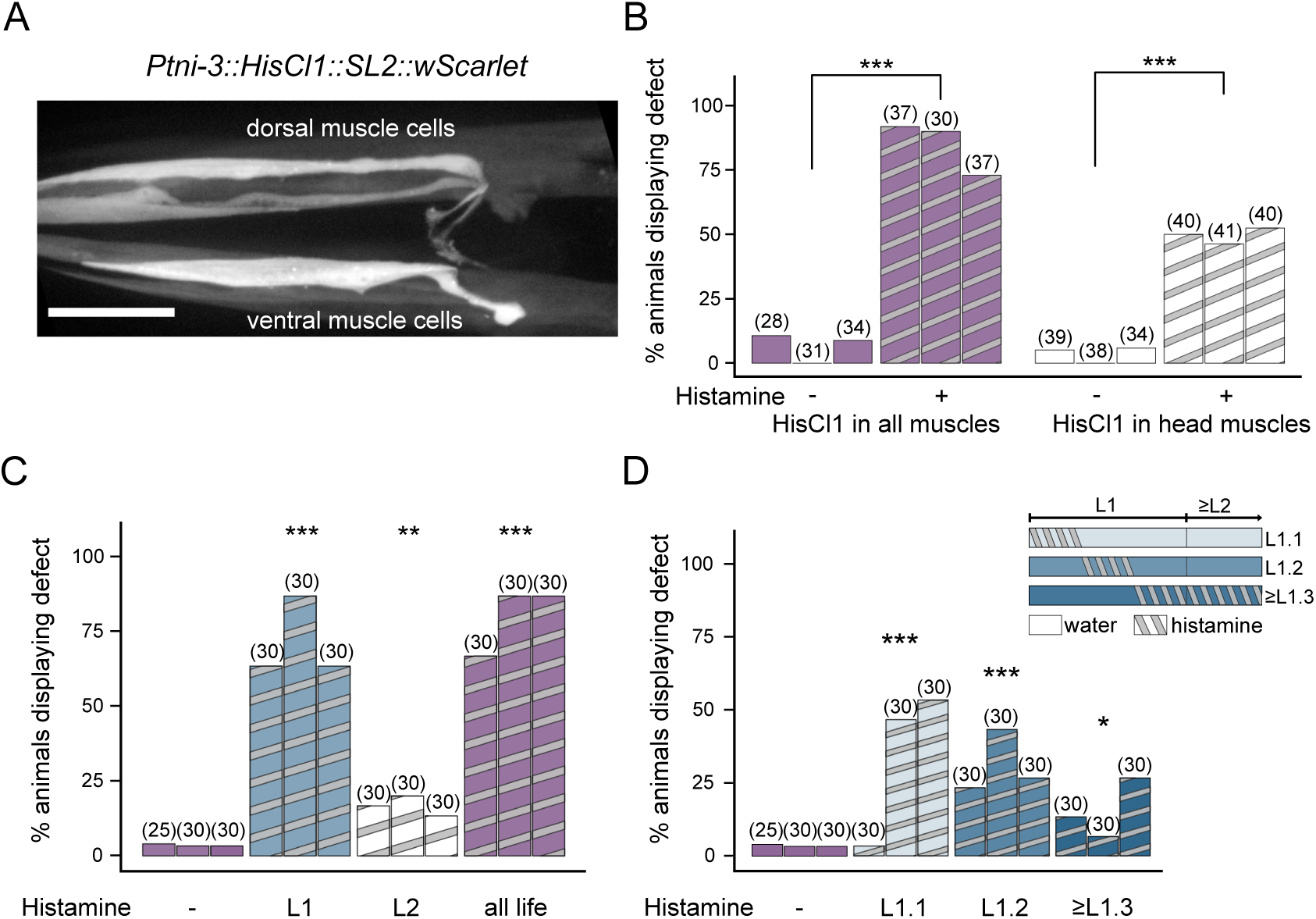
SAB axonal overgrowth is triggered by inhibition of muscle activity at early developmental stages and can be induced by silencing head muscles alone. (**A**) Confocal image of the head of an animal carrying the multi-copy integrated transgene *Ptni-3::HisCl1::SL2::wScarlet*. Nose is on the left, pharyngeal bulbs on the right. Scale bar is 20 µm. (**B**) Percentage of animals with SAB defects when HisCl1 is expressed in all muscles or only head muscle, following water (-) or histamine (+) treatment. (**C**) Percentage of control animals (*Pmyo-3::HisCl1::SL2::tagBFP*; *Punc-4::GFP*) with SAB defects, after water treatment (none), or histamine exposure restricted to the L1 stage (L1), starting from the L2 stage (L2) or throughout life (all life). (**D**) Percentage of control animals with SAB defects, after water treatment (none) or histamine exposure during the first third of L1 stage alone (L1.1), the second third of L1 stage alone (L1.2) or starting from the last third of L1 stage (≥L1.3). Data in (C) and (D) are from the same experimental batch and have been plotted separately for visual clarity. Therefore, the water-treated controls (-) are identical in both panels. Bars represent independent replicates (ordered left to right by experiment date, matched across cohorts). Numbers above bars indicate the number of animals scored. Statistical significance was determined by CMH test against water-treated controls *: p<0.05, **: p<0.01, ***: p<0.001.

### The critical time window for activity-dependent axon termination occurs during the L1 stage

Previous work by Zhao and Nonet showed that reduced synaptic activity must occur before the third larval stage (L3) to induce SAB axonal overgrowth (Zhao and Nonet, 2000). To define the critical window in our experimental paradigm, we inhibited muscle activity during specific larval stages (Figure 2C). Continuous histamine exposure throughout larval development induced strong SAB axonal overgrowth, whereas untreated controls showed minimal defects. Histamine treatment limited to the first larval stage (L1) produced a strong effect, comparable to continuous exposure. By contrast, treatment limited to the second larval stage (L2) resulted in limited overgrowth (Figure 2C). To further refine the critical period, we restricted histamine exposure to three distinct sub-phases of the L1 stage. Histamine treatment during the early, middle, or late L1 stage alone was sufficient to induce moderate SAB overgrowth (Figure 2D). These findings suggest that SAB axonal termination is sensitive to muscle activity throughout the entire L1 larval stage, and mostly during this developmental window.

### Neuronal voltage-gated calcium channels are involved in SAB axonal overgrowth

L-type voltage-gated calcium channels (VGCCs) negatively regulate axon termination in two mechanosensory neurons of *C. elegans* (Buddell et al., 2019; Buddell and Quinn, 2021). To investigate whether VGCCs also play a role in SAB axonal overgrowth triggered by reduced muscle activity, we examined SAB axons in mutants carrying loss-of-function mutations in the genes encoding the three pore-forming VGCC subunits: EGL-19 (L-type) (Lee et al., 1997), UNC-2 (P/Q type) (Schafer and Kenyon, 1995) and CCA-1 (T-type) (Steger et al., 2005); as well as UNC-36, an auxiliary α2/δ subunit shared by P/Q- and L-type VGCCs (Frøkjaer-Jensen et al., 2006; Lainé et al., 2011; Saheki and Bargmann, 2009).

Upon histamine exposure, *egl-19* hypomorphic mutants displayed a strong suppression of SAB overgrowth induced by muscle activity (Figure 3A, S1). Similarly, overgrowth was also significantly reduced in *unc-2* and *unc-36* mutants (Figure 3A). In contrast, *cca-1* mutants showed no suppression.

**Figure 3.**
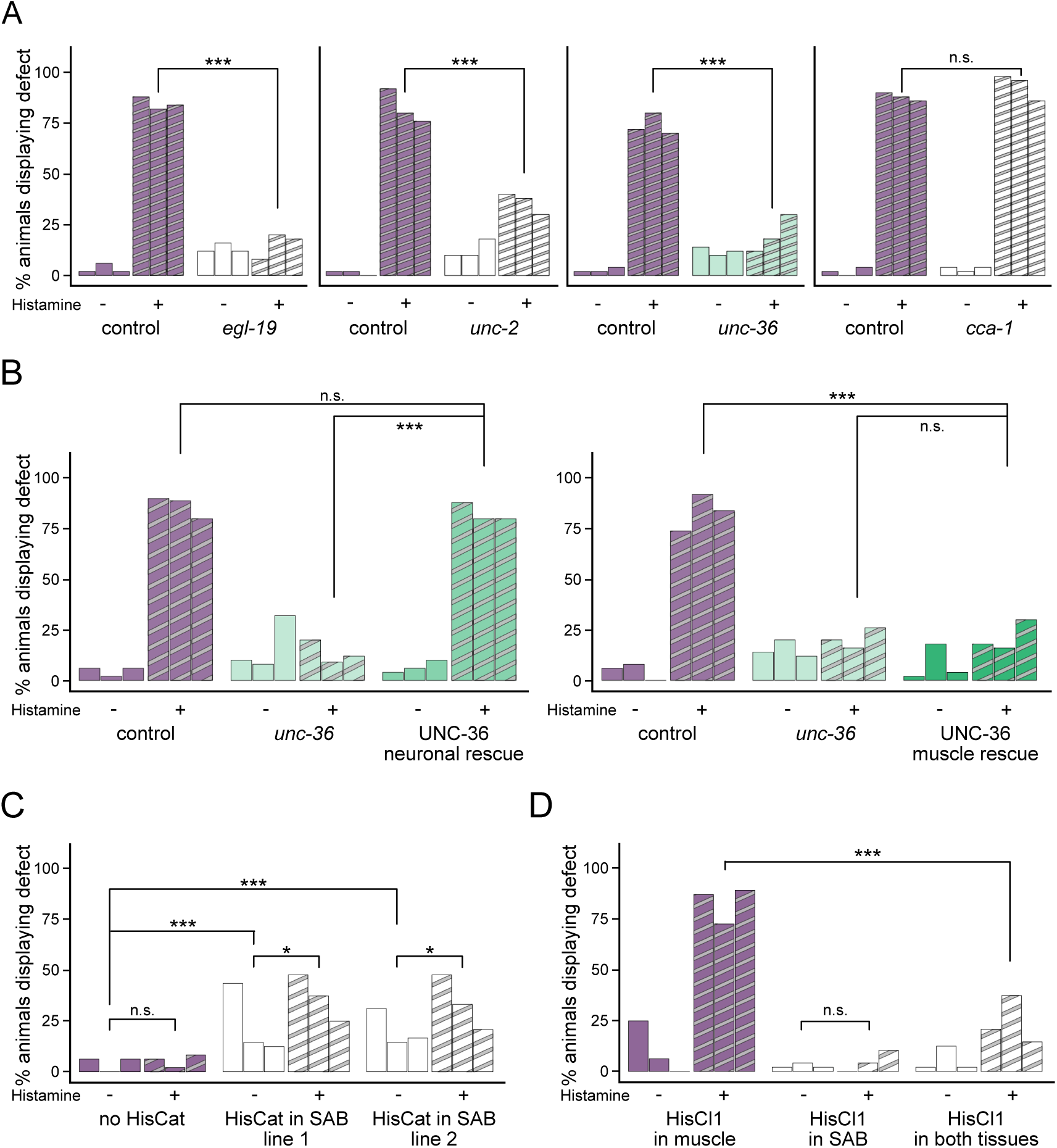
Activity-dependent SAB axonal overgrowth depends on neuronal VGCC. (**A**) Percentage of control animals and *egl-19(n582)*, *unc-2(e55)*, *unc-36(e251)* and *cca-1(ad1650)* mutants with SAB defects, after treatment with water (-) and 10 mM histamine (+). For each genotype, condition and replicate N=50. (**B**) Percentage of animals with SAB defects in control, *unc-36(e251)* single mutants and *unc-36(e251)* animals carrying a single copy integrated transgene *Punc-4::GFP-unc-36* (UNC-36 neuronal rescue, left panel) or *Pmyo-3::GFP-unc-36* (UNC-36 muscle rescue, right panel). N=50, except in the left panel for: replicate 2 of control with histamine N=45, replicates 2 and 3 of *unc-36* with histamine, N=22 and 25, respectively. Animals were treated by water (-) or 10 mM histamine (+). (**C**) Percentage of animals with SAB defects in *Punc-4::mNeonGreen* animals carrying no supplemental transgene (no HisCat) or a multicopy integrated transgene expressing the histamine-gated cation channel HisCat in SAB (HisCat in SAB, line 1 and 2). N=50. Animals were treated by water (-) or 30 mM histamine (+). (**D**) Percentage of animals with SAB defects in *Punc-4::mNeonGreen* animals carrying a single-copy integrated transgene expressing HisCl1 channel in muscle (*Pmyo-3::HisCl1::SL2::tagBFP*), SAB neurons (*Punc-4::HisCl1::SL2::tagBFP*) or in both tissues (*Pmyo-3::HisCl1::SL2::tagBFP; Punc-4::HisCl1::SL2::tagBFP*). N=50. Animals were treated by water (-) or 10 mM histamine (+). In all panels, bars represent independent replicates (ordered left to right by experiment date, matched across cohorts). Statistical significance was determined by CMH tests, n.s.: non significant, *: p<0.05, ***: p<0.001.

To identify the tissue in which VGCCs act, we performed tissue-specific rescue experiments in an *unc-36* null mutant background. Re-expression of *unc-36* in neurons fully restored the overgrowth phenotype (Figure 3B). In contrast, re-expression of *unc-36* in muscle had no significant effect (Figure 3B). These findings demonstrate that VGCCs function cell-autonomously in SAB neurons to mediate axonal overgrowth in response to reduced muscle activity.

### Motor neuron hyperactivity mediates axonal overgrowth

Based on our results, we hypothesized that inhibition of muscle activity triggers hyperactivity in SAB motor neurons, which in turn drives axonal overgrowth. In this model, reducing VGCC activity in neurons counteracts SAB hyperactivity and suppresses the overgrowth phenotype.

The first prediction of this model is that increasing activity in SAB neurons should induce axonal overgrowth. To test this, we expressed a modified version of the HisCl1 channel, named HisCat, in SAB neurons. HisCat is a histamine-gated cation channel that depolarizes neurons upon activation, thereby inducing hyperactivity (gift from M. Alkema, unpublished). We generated two independent extra-chromosomal transgenic lines expressing the HisCat channel specifically in SAB motor neurons. Upon histamine treatment, both lines displayed a significantly higher rate of SAB axonal overgrowth compared to controls expressing only mNeonGreen in SAB (Figure 3C). Interestingly, even in the absence of histamine, HisCat-expressing animals exhibited elevated incidence of axonal defects relative to animals expressing only mNeonGreen, suggesting that the transgene may be active without histamine treatment.

The second prediction of our model is that decreasing SAB activity in the context of muscle silencing should suppress SAB axonal overgrowth. To test this, we expressed the HisCl1 channel in SAB neurons of animals already expressing HisCl1 in muscle. Inhibiting SAB activity alone had no effect while decreasing muscle activity induced robust SAB overgrowth (80% of animals affected) (Figure 3D). By contrast, simultaneous reduction of both muscle and SAB neuron activity led to a strong suppression of SAB axonal overgrowth. These results indicate that inhibition of muscle activity promotes axonal overgrowth by increasing the intrinsic excitability of SAB motor neurons.

### Neuropeptide signaling modulates activity-dependent SAB axonal termination

We then sought to explore the molecular nature of the signal that muscles send to SAB neurons. Neuropeptides have been proposed as potential retrograde signals in both early and recent studies (Doi and Iwasaki, 2002; Rennich et al., 2025). We tested this by assessing SAB axonal morphology following muscle inhibition in mutants with impaired neuropeptide production. The *egl-3* gene encodes a proprotein convertase essential for processing most neuropeptides (Husson et al., 2006). Upon muscle inhibition, *egl-3* mutants exhibited a two-fold reduction in SAB axonal overgrowth compared to control animals (Figure 4A).

**Figure 4.**
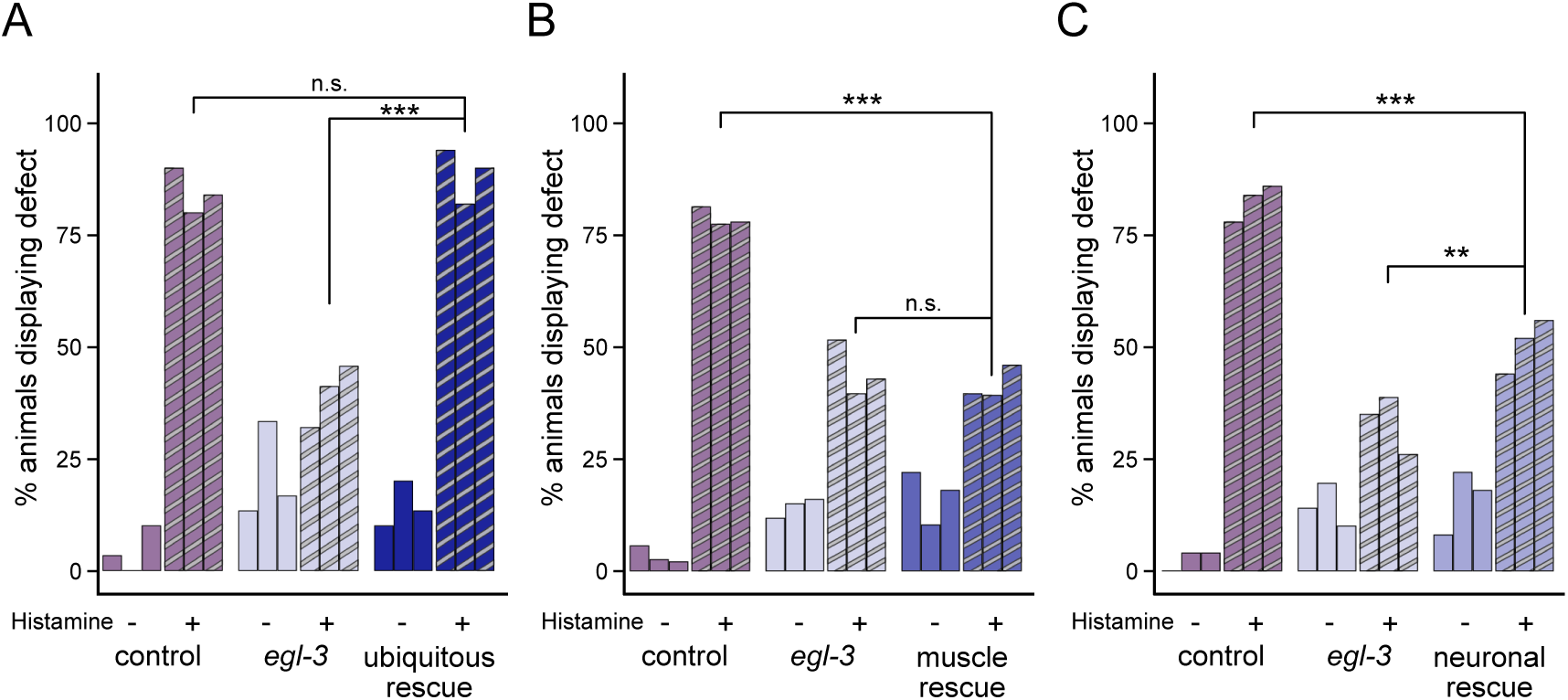
Activity-dependent SAB axonal overgrowth depends on neuropeptide signaling. (**A**) Percentage of animals with SAB defects in control, *egl-3(gk238)* single mutant and *egl-3(gk238)* animals carrying a single copy integrated transgene *Peft-3::egl-3(cDNA)::SL2::wScarlet* (ubiquitous rescue). For genotypes treated with water, N=30. For genotypes treated with histamine N=50, except in EGL-3 ubiquitous rescue where N=50, 22 and 25. (**B**) Percentage of animals with SAB defects in control, *egl-3(gk238)* single mutant and *egl-3(gk238)* animals carrying a single copy integrated transgene *Pmyo-3::egl-3(cDNA)::SL2::wScarlet* (muscle rescue). For control treated with water N=36, 40 and 50, for control treated with histamine N=43, 40 and 50, for *egl-3* treated with water N=51, 40 and 50, for *egl-3* treated with histamine N=31, 48 and 40, for EGL-3 muscle rescue treated with water N=50, 39 and 50, for muscle rescue treated with histamine N=48, 51, 50. (**C**) Percentage of animals with SAB defects in control, *egl-3(gk238)* single mutant and *egl-3(gk238)* animals carrying a single copy integrated transgene *Prgef-1::egl-3(cDNA)::SL2::wScarlet* (neuronal rescue). N=50 for all, except in water-treated *egl-3* where N=46 for replicate 2 and histamine-treated egl-3 with N=40, 49 and 50. In all panels, animals were treated by water (-) or 10 mM histamine (+). Bars represent independent replicates (ordered left to right by experiment date, matched across cohorts). Statistical significance was determined by CMH tests, n.s.: non significant, *: p<0.05, **: p<0.01.

To determine in which tissue *egl-3* functions to regulate this phenotype, we re-expressed the *egl-3* gene in a null mutant background using several tissue-specific promoters. Ubiquitous re-expression of *egl-3* restored overgrowth to control levels (Figure 4A). By contrast, re-expressing *egl-3* in muscle alone failed to rescue the phenotype, with a lower occurrence of overgrowth resembling that observed in *egl-3* mutants rather than in control animals (Figure 4B). Pan-neuronal expression of *egl-3* partially rescued the phenotype, although it was still significantly lower than that of control animals (Figure 4C). These data suggest that EGL-3 plays a role in neurons, but not in muscles, in the activity-regulated overgrowth of SAB axons.

To identify the neuropeptide genes involved in SAB axonal termination, we used an unbiased transcriptomic approach. We analyzed gene expression changes following muscle inhibition in control animals collected during the critical L1 period. A naïve differential test yielded 1,602 differentially expressed genes (777 upregulated and 825 downregulated). To further characterize these genes, we performed Gene Ontology enrichment analysis and found significant associations with multiple processes related to larval development, including muscle development, ribosome components, or molting cycle (Figure 5A). This pattern suggests that the observed differences in gene expression may in part reflect a developmental delay caused by histamine-induced muscle inhibition. To investigate this possibility, we monitored the timing of L1 to L2 molt by measuring the period of animal quiescence. Histamine treatment delayed molting of control animals by nearly four hours (Figure 5B,C). To correct for developmental delay as a confounding factor, we used RAPToR—a computational tool that estimates and corrects the developmental age of a sample based on its transcriptome (Bulteau and Francesconi, 2022). Estimates of sample age by RAPToR confirmed the developmental delay following histamine treatment (Figure 5D). After correcting for age differences using both the “larval” and “larval + young adult” references, we intersected the resulting gene sets and retained only genes differentially expressed in both corrected tests as well as the uncorrected test. This yielded 469 differentially expressed genes (366 upregulated and 103 downregulated). Gene Ontology analysis no longer showed development-associated terms; instead, the top enriched category was “neuropeptide signaling pathway”, containing 24 genes, of which 12 encoded neuropeptide genes (Figure 5E, Table 1).

**Figure 5.**
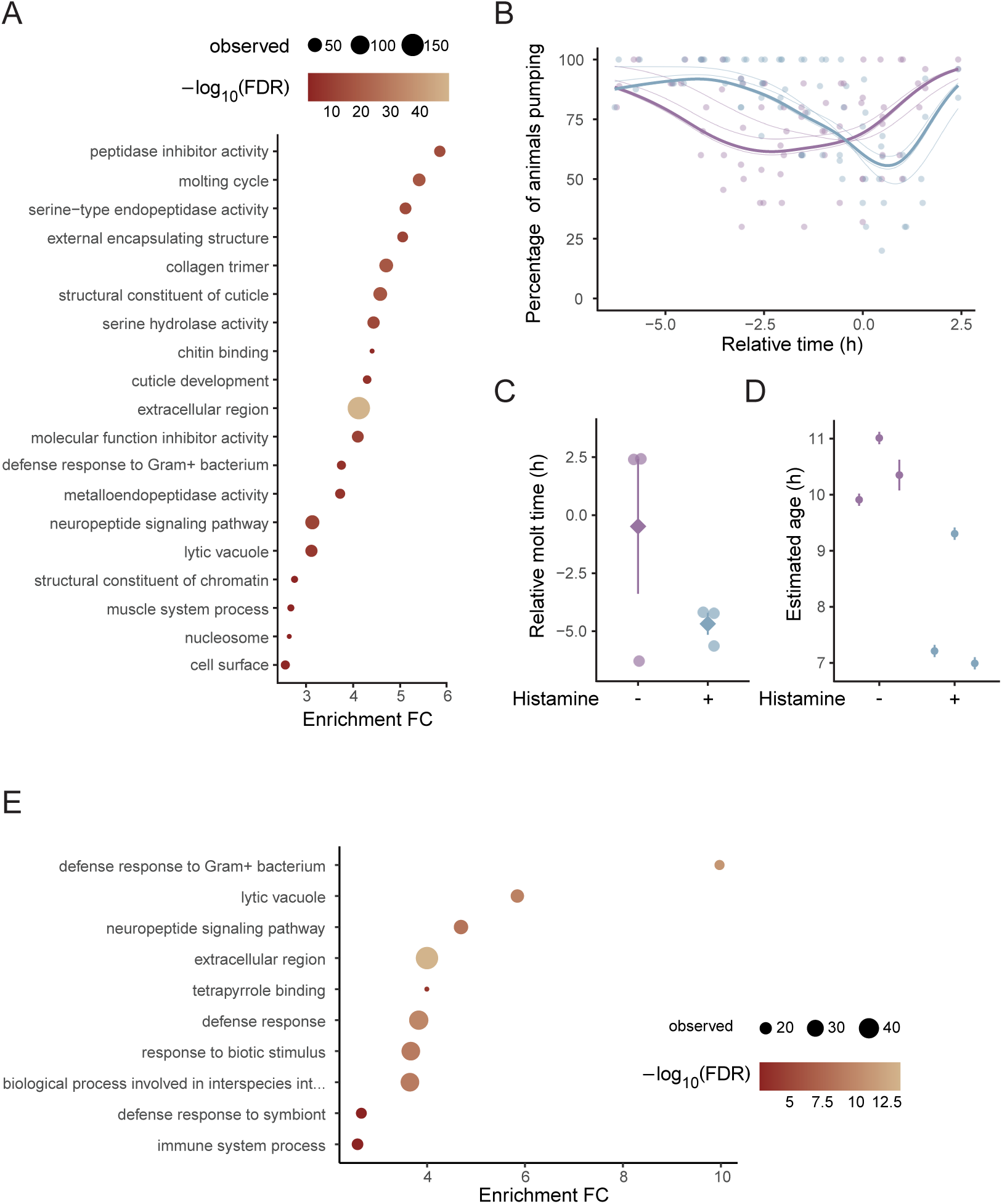
Transcriptomic analysis after age-correction by RAPToR reveals an increase in neuropeptide synthesis. (**A**) Gene Ontology terms from naïve Differential Expression. (**B**) Percentage of control worms pumping food around the L2 to L3 molt. Points indicate the percentage at individual timepoints, the thin lines are GAMM fits of individual replicates, the thick lines the population-level fits. Time is indicated in hours relative to timing of molt for non-transgenic animals. (**C**) Time of molt, i.e. minimum of curve from B. (**D**) Age of sequenced samples estimated with RAPToR using the larval reference. (**E**) GO terms after correction. All animals were treated with histamine (purple) or water (blue).

To assess the role of neuropeptides in SAB activity-dependent axonal overgrowth, we selected four neuropeptide genes – *flp-18*, *nlp-12*, *nlp-11* and *nlp-21 –* and analyzed the corresponding mutants. *flp-18* and *nlp-12* mutants showed a reduction in SAB axonal overgrowth in response to muscle inhibition compared to controls (Figure 6A, B). By contrast, *nlp-11* and *nlp-21* mutants displayed axonal defects at a rate similar to that of control animals (Figure 6C, D). To determine whether FLP-18 and NLP-12 act in parallel or within the same signaling pathway, we generated a *nlp-12; flp-18* double mutant and quantified the incidence of axonal overgrowth. Upon muscle inhibition, the double mutants exhibited a rate of animals with axonal defects significantly lower than that observed in *nlp-12* single mutants. These results indicate that both FLP-18 and NLP-12 modulate SAB axonal overgrowth in response to reduced muscle activity, and that they likely function through partially overlapping yet distinct signaling pathways.

**Figure 6.**
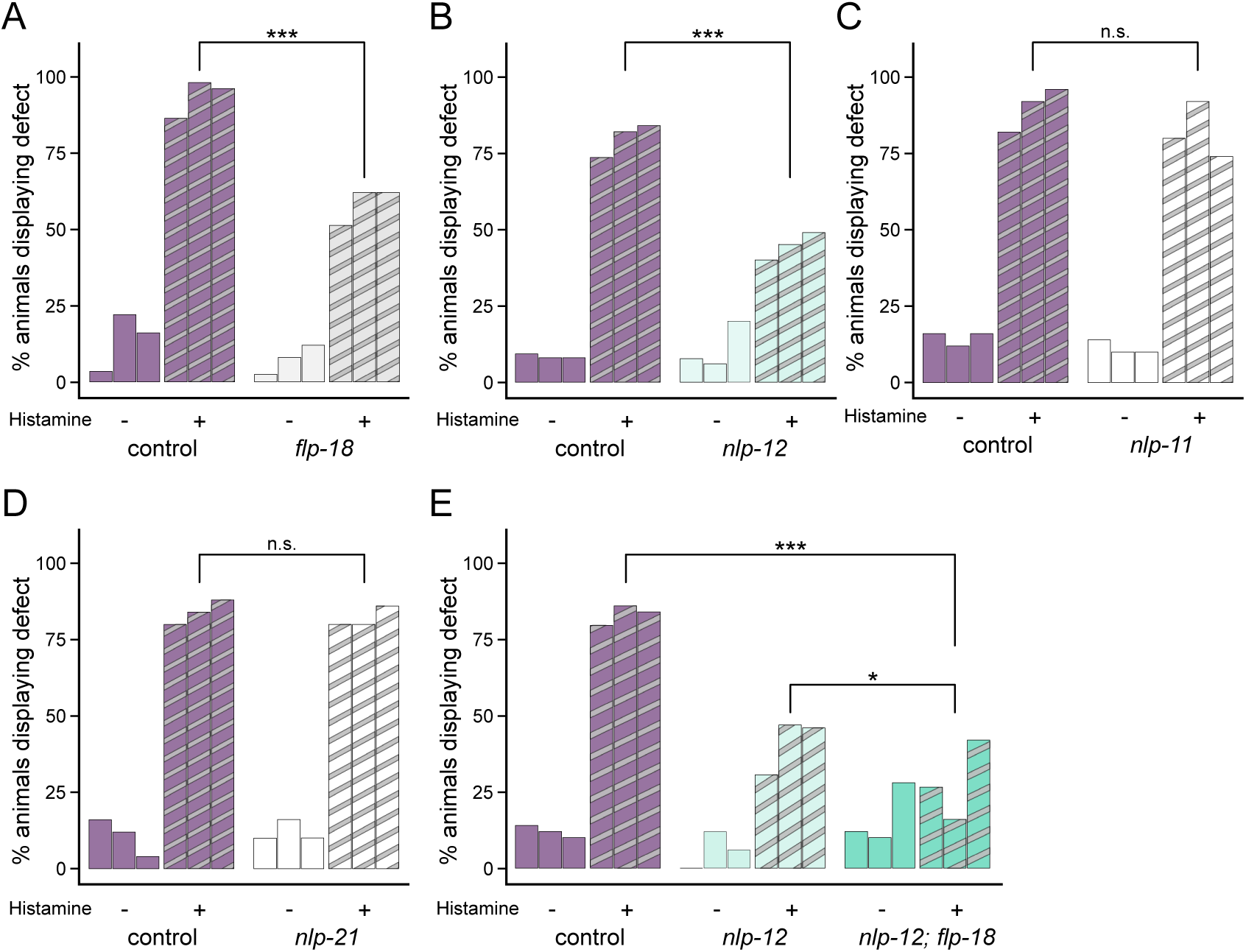
*flp-18* and *nlp-12* mutations suppress activity-induced SAB axonal defects. (**A**) Percentages of control animals and *flp-18(tm2179)* mutants with SAB defects, after treatment with water (-) or 10 mM histamine (+). N=50, except in replicate 1, where N=58, 51, 41 and 42 for water-treated controls, histamine-treated controls, water-treated *flp-18* and histamine-treated *flp-18*, respectively. (**B**) Percentages of control animals and *nlp-12(ok335)*mutants with SAB defects, after treatment with water (-) or 10 mM histamine (+). N=50, except in replicate 1 where N=54, 53, 52 and 55 for water-treated controls, histamine-treated controls, water-treated *nlp-12* and histamine-treated *nlp-12*, respectively; in replicate 2 where N=51 for histamine-treated *nlp-12*; and in replicate 3 where N=49 for histamine-treated *nlp-12*. (**C**) Percentages of control animals and *nlp-11(yum480)* mutants with SAB defects, after treatment with water (-) or 10 mM histamine (+). N=50. (**D**) Percentages of control animals and *nlp-21(yum445)* mutants with SAB defects, after treatment with water (-) or 10 mM histamine (+). N=50. (**E**) Percentages of control animals, *nlp-12(ok335)* mutants and *nlp-12(ok335); flp-18(tm2179)* double mutants, after treatment with water (-) or 10 mM histamine (+). N=50, except in replicate 1 where N=49 for histamine-treated animals and N=40 for water-treated controls, in replicate 2 where N=51 for histamine-treated controls and N=48 for histamine treated double mutants. In all panels, bars represent independent replicates (ordered left to right by experiment date, matched across cohorts). Statistical significance was determined by CMH tests, n.s.: non significant, *: p<0.05, ***: p<0.001.

## Discussion

In this study, we identify a retrograde signaling mechanism through which head muscle activity constrains SAB motor axon outgrowth during early development. We find that this effect is temporally restricted to the L1 stage, depends on neuronal calcium influx through L-type VGCCs, and requires the neuropeptides *flp-18* and *nlp-12*.

### Activity-dependent retrograde signaling at the neuromuscular junction during a critical period

Our findings reveal striking parallels with activity-dependent retrograde mechanisms described at neuromuscular junctions in other organisms. In mammals, motor axons frequently sprout following denervation or inactivity, yet the nature of the retrograde signals driving this response remains poorly understood (Brown et al., 1981; English, 2003; Slater, 2017). During development, muscle-derived neurotrophic factors, such as brain-derived neurotrophic factor (BDNF), glia cell line-derived neurotrophic factor (GDNF), and the neurotrophins NT-3 and NT-4, have been implicated as activity-dependent retrograde cues. These factors prevent neuronal degeneration and promote axon extension, in part by acting through presynaptic calcium signaling (Funakoshi et al., 1995; Keller-Peck et al., 2001; Koliatsos et al., 1993; Stanga et al., 2021; Zhan et al., 2003).

In *C. elegans*, we find that SAB axons exhibit a comparable form of activity-dependent overgrowth. A similar phenotype has been reported for the DVB motoneuron when its target enteric muscles are inhibited (Hobert et al., 1999; Loria et al., 2003). These observations suggest that activity-dependent retrograde signaling constitutes a general feature of neuromuscular junction development. Importantly, SAB overgrowth occurs only when muscle is silenced during the L1 stage, revealing the existence of a defined critical period. Comparable critical windows of plasticity are well documented in the visual system (Hubel and Wiesel, 1970; Kutsarova et al., 2016). More broadly, experience-dependent plasticity has been described in the rodent barrel cortex (Jamann et al., 2018) and in *C. elegans* sensory neurons (Peckol et al., 1999). Our results therefore suggest that SAB motor neurons, like sensory circuits, can be constrained by temporally defined, activity-dependent processes.

### Calcium and excitability as brakes on axon growth

We further show that neuronal calcium influx is required to suppress SAB overgrowth. This aligns with evidence from other remodeling contexts in *C. elegans*, where calcium entry through the DEG/ENaC channel UNC-8 drives disassembly of presynaptic structures during DD remodeling (Cuentas-Condori et al., 2023) (Miller-Flemming et al., 2015; Cuentas-Condori et al., 2023). Studies on the PLM and ALM mechanosensory neurons have demonstrated that activity of the L-type VGCC *egl-19* and the P/Q-type *unc-2* regulates axon termination (Buddell et al., 2019; Buddell and Quinn, 2021; Ghosh-Roy et al., 2010). Similarly, at the *Drosophila* neuromuscular junction, calcium-dependent activation of CaMKII shapes presynaptic morphology (Nesler et al., 2016). In the mammalian cortex, calcium entry through L-type VGCCs limits axon elongation and stabilizes existing projections (Hutchins and Kalil, 2008; Tang et al., 2003). Even in contexts of axon competition, neurotrophin signaling requires local depolarization and calcium influx to promote axon stabilization via CaMKII (Singh and Miller, 2005). Taken together, these findings support the view that calcium influx functions as a conserved molecular brake, terminating axon extension once appropriate contacts are established.

### Neuropeptides as retrograde modulators

Our genetic analysis implicates the neuropeptides *flp-18* and *nlp-12* in the suppression of SAB overgrowth. Neuropeptides are widely recognized as key neuromodulators in *C. elegans*, influencing circuit dynamics and behavior (Zhen and Samuel, 2015). In some cases, they participate in retrograde pathways: for example, postsynaptic expression of AEX-1 and AEX-5 in muscle and intestine regulates presynaptic activity, consistent with peptide-mediated retrograde signaling (Doi and Iwasaki, 2002). Our findings extend this principle by identifying *flp-18* and *nlp-12* as candidate mediators of activity-dependent suppression of SAB axon growth.

*flp-18* expression is activity-dependent and plays a central role in the homeostatic regulation of locomotor circuits. Overexcitation of cholinergic motor neurons leads to increased expression of *flp-18* (McCulloch et al., 2020; Stawicki et al., 2013). FLP-18 exerts a homeostatic effect on the locomotor circuit by enhancing inhibitory GABAergic transmission at the neuromuscular junction (Stawicki et al., 2013). FLP-18 can also act through the G-protein-coupled receptor NPR-4 to modulate calcium dynamics in AVA interneurons (Bhardwaj et al., 2018). Together with NPR-4 and NPR-5, FLP-18 forms “pervasive” signaling networks that involve multiple releasing neurons and diverse downstream targets, including muscles (Ripoll-Sánchez et al., 2023; Stawicki et al., 2013).

*nlp-12*, the *C. elegans* homolog of cholecystokinin, is produced primarily by the stretch-sensitive DVA interneuron (Bhattacharya et al., 2014; Hu et al., 2011; Li et al., 2006). NLP-12 release is activity-dependent and potentiates cholinergic transmission to muscles, forming a positive feedback loop (Bhattacharya et al., 2014; Hu et al., 2015, 2011). DVA integrates multiple inputs, including dopaminergic signals, to modulate motor behavior during local food-searching strategies (Bhattacharya and Francis, 2015). Acting through CKR-1 and CKR-2, NLP-12 serves as the core of a “broadcasting” network to reach multiple downstream targets—the previously described “local” architecture of the CKR-2 network likely reflects a threshold rather than a fundamentally restricted network (Ripoll-Sánchez et al., 2023).

As SAB overgrowth is not induced by generalized paralysis from myosin mutation but is triggered by selective inhibition of head muscles, the relevant retrograde signal is unlikely to arise solely from stretch-dependent DVA activation. Importantly, since *egl-3* is required in neurons rather than muscle, the direct retrograde signal from muscle remains unidentified. Our data instead suggest that *flp-18* and *nlp-12* act as downstream circuit modulators that integrate muscle activity with presynaptic excitability to regulate axon growth.

### Characteristics distinguishing the SAB neurons

Interestingly, other motor neurons, including the VA and DA neurons of the ventral cord, do not display overgrowth following inhibition of all body-wall muscles, highlighting a unique plasticity of the SAB neurons (Zhao and Nonet, 2000). This specificity may reflect functional, anatomical, and molecular differences. Functionally, the SABs innervate head muscles that control subtle snout movements and require a specific neuronal architecture (Emmons, 2024; White, 2018). NLP-12 differentially fine-tunes the head and body muscle activities to match behavioral goals (Ramachandran et al., 2021). Anatomically, SAB axons extend within the head neuropil, which offers greater spatial freedom for neurite outgrowth compared with the tightly bundled axons of the ventral nerve cord (White et al., 1986, 1976). Molecularly, SABs display preferential expression of glutamate receptors (*glr-4* and *glr-5*) relative to A-type motor neurons of the ventral cord (Brockie et al., 2001; Taylor et al., 2021) that may facilitate crosstalk with head circuits important for foraging. Together, these features may underlie the distinct plasticity of SAB compared with ventral cord motor neurons.

### Perspectives

Our findings implicate FLP-18 and NLP-12 as candidate mediators of activity-dependent control of SAB axon growth, but several key questions remain. Future work should identify the relevant neuropeptide receptors and dissect their contributions through tissue-specific mutations and rescue experiments. Moreover, the identity of the retrograde signal from muscle remains unknown. Finally, determining how neuromodulators such as NLP-12 integrate head muscle activity with behavioral state will shed light on how motor circuits balance stability with plasticity.

## Methods

### *C. elegans* genetics and molecular biology

Animals were maintained and bred at 20°C on nematode growth medium (NGM) plates seeded with *Escherichia coli* OP50, according to standard protocols (Brenner, 1974). All strains used in this study were derived from the wild-type Bristol N2 strain (Table S1). Plasmids were generated by Gibson assembly (Gibson, 2011) and all open reading frames were verified by Sanger sequencing.

Transgenesis was performed by microinjections into the gonads of adult hermaphrodites, following standard protocols (Mello and Fire, 1995). Single-copy insertions were generated using the miniMos insertion method (Frøkjær-Jensen et al., 2014), while multi-copy insertions were obtained either by UV-irradiation (Mariol et al., 2013) or the MosTi method (El Mouridi et al., 2022). Selection markers included hygromycin resistance for *krSi16* and neomycin resistance for *krSi24*.

### Histamine treatment

To inhibit SAB or muscle activity, histamine-gated chloride channels (HisCl1) were expressed under the control of tissue-specific promoters (Pokala et al., 2014). For each experiment and each genotype, around 20 gravid animals were placed on NGM plates to lay eggs for 2-5 hours and then removed. Subsequently, 300 µL of water (control plates) or 300 µL of histamine (histamine plates) 1/30 M was added to 10 mg NGM plates. Progeny were allowed to develop for three days at 20°C. To increase SAB electrical activity, we used a histamine-gated cationic channel (HisCat, kind gift of Mark Alkema, UMass Chan Medical School), activated by 30 mM of histamine. Histamine (Sigma-Aldrich) was stored as 1 M aqueous stock solution.

### Microscopy and SAB scoring

SAB axonal morphology was visualized by expressing mNeonGreen under a truncated *unc-4* promoter (promC), which drives strong expression in SABs and weak expression in other neurons (Kratsios et al., 2015). After water or histamine treatment, animals were mounted on dry pads (2% agarose in water, dry overnight) and immobilized with 4% polyethylene beads (Polysciences, 0.1 µm).

SAB axonal defects were scored at 60x magnification using either a Zeiss Axioskop upright microscope equipped with a Photometrics CoolSNAP HQ^2^ CCD camera or a Nikon Eclipse Ti2 inverted microscope with a Photometrics Prime BSI CMOS camera. Animals were scored as defective if at least one of their four SAB axons exhibited a loop or a sprout. The reported n corresponds to the number of individual worms analyzed. Three replicates were performed per condition/genotype, except in Figure S1 (two replicates).

Postsynaptic bouton density was quantified at 60x using an Olympus IX83 inverted microscope equipped with a Yokogawa confocal scanner unit spinning-disk scan head and an Andor iXon Ultra 888 EMCCD camera. Synapses were counted along clearly visible SAB axons and normalized to the length of the portion observed.

Epifluorescence and confocal images were processed and analyzed using the open-source image processing software Fiji ImageJ (version 2.3.0/1.53q).

### RNA-Seq

To obtain synchronized populations, around 50 gravid control (*Pmyo-3::HisCl1::SL2::tagBFP*; *Punc-4::GFP*) adults were allowed to lay eggs for two hours before removal, yielding 300-600 progeny per plate. Plates were supplemented with histamine or water (control). After two-thirds of the L1 stage, larvae were collected in cold M9 buffer. A subset of individuals was isolated and kept for phenotypic validation, while the remainder was resuspended in QIAzol buffer and stored at −80°C. After three days, the isolated individuals were examined, and only samples with a high occurrence of SAB defects were further processed. Cuticles were disrupted by five freeze-thaw cycles in liquid nitrogen, and RNA was isolated using the miRNeasy Mini kit (Qiagen, catalog number 217004), including the optional DNase I treatment. Three biological replicates were collected per condition. Sequencing was performed on an Illumina HiSeq 4000.

Reads were pseudo-aligned to the WormBase WS298 transcriptome (Sternberg et al., 2024) using kallisto 0.44.0 (Bray et al., 2016). After filtering, 15,962 genes with at least 10 reads in at least 3 samples were retained. Differential expression analysis was done using DESeq2 v1.44.0 (Love et al., 2014), with genes considered differentially expressed at FDR < 0.05 and log fold change > 0. Gene Ontology enrichment was performed against WormBase annotations (Angeles-Albores et al., 2018) using all 15,944 tested genes as a background list. Enriched terms were defined at FDR < 0.05, enrichment fold change > 2.5, and at least 10 genes observed.

Developmental stage correction was performed using RAPToR with the “Cel_larval” and “Cel_larv_YA” references (Bulteau and Francesconi, 2022). Briefly, the interpolated reference data was transformed into artificial counts which was analyzed along the experimental data, including the developmental time as a covariate.

Molting delays were estimated using synchronized populations of control animals obtained by timed egg-lays. From 28-37 hours post egg-lay at 20°C (L2-L3 molt), groups of 10-30 animals were scored for feeding behavior at 30 min intervals, to determine the proportion of animals actively feeding or quiescent (Singh and Sulston, 1978). Experiments were performed in triplicate, including untreated wild-type animals (not shown). Timing of minimal pharyngeal pumping in untreated wild type was used as a reference; the other time points were given in time relative to this reference. We then fitted a generalized additive mixed model (GAMM) with a binomial distribution to model the proportion of non-quiescent animals using cubic regression splines for time and allowing random intercepts and slopes per replicate. Minimal pumping times were extracted from smoothed curves and compared between water and histamine-treated control animals using two-sided Student t-tests.

### Data representation and statistics

For SAB axonal defect assays, three replicates were performed per genotype, except in Figure S1 (two replicates). Replicates were plotted individually as bar graphs, to better illustrate biological variability. Bars were ordered from left to right by date, such that corresponding positions represent matched replicates.

Penetrance of SAB defect was assessed with the Cochran–Mantel–Haenszel (CMH) test (Cochran, 1954; Mantel and Haenszel, 1959), a modified *χ*^2^ test that stratifies data by replicate (date) to account for day-to-day variability. For each replicate, 2 x 2 contingency tables were constructed based on binary outcome (normal SAB axons *versus* defective SAB axons) and predictors (genotype or histamine treatment).

For postsynaptic bouton density assays, data were represented as violin plots, and significance was assessed by Mann–Whitney tests.

All statistical analyses were performed in R. Significance was reported as: n.s.: not significant, *: p < 0.05, **: p < 0.01, ***: p < 0.001. Raw data and p-values of the statistical tests are provided in Supplementary Information.

## Acknowledgments

We thank Driss Laabid for technical assistance, and Thomas Boulin, Paschalis Kratsios and Isabel Beets for scientific discussions. We are grateful to Sonia El Mouridi and Christian Frøkjær-Jensen for providing MosTI reagents, Mark Alkema for the HisCat plasmid, and Mario de Bono, Changchun Chen and Oliver Hobert for sharing strains. We acknowledge the CIQLE Imaging facility, member of the national infrastructure France-BioImaging supported by the French National Research Agency (ANR-24-INBS-0005 FBI BIOGEN) for access to equipment and GenomEast platform, a member of the “France Génomique” consortium (ANR-10-INBS-0009). Most strains were provided by the *Caenorhabditis* Genetics Center, which is funded by NIH Office of Research Infrastructure Programs (P40 OD010440).

