## Supplementary information for "Axon termination of the SAB motor neurons in *C. elegans* depends on pre- and postsynaptic activity"

**Figure S1. *egl-19(ad1006)* mutation suppresses activity-induced axonal defect of SAB.**
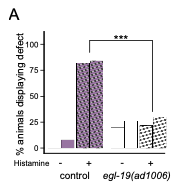

Percentage of control animals and *egl-19(ad1006)* mutants with SAB defects. Bars represent two independent replicates. For each genotype and each condition, N=50.

**Table S1. List of the strains used in this study.**

| Figure | Name | Genotype | Transgenesis |
| --- | --- | --- | --- |
| 1B-D, F ; 2B-D ; 3A,B,D ; 4A-C ; 5 ; 6A-G | EN3940 | *krSi16[Pmyo-3::HisCl1::SL2::tagBFP]I* ; *krSi24[Punc-4::mNeonGreen]* | miniMos single copy insertion in EN3831 |
| 1C ; 3C | EN3831 | *krSi24[Punc-4::mNeonGreen]* | miniMos single copy insertion |
| 1D | EN2383 | *unc-29[kr208-tag-RFP-T]I* ; *jsIs42[Punc-4::snb-1-GFP]X* |  |
| 1E | EN3939 | *unc-29[kr208-tag-RFP-T]I* *krSi16[Pmyo-3::HisCl1::SL2::tagBFP]I* ; *krSi24[Punc-4::mNeonGreen]* |  |
| 1F | EN7133 | *unc-54[e190]I* *krSi16[Pmyo-3::HisCl1::SL2::tagBFP]I* ; *krSi24[Punc-4::mNeonGreen]* |  |
| 2A-B | EN7922 | *krIs80[Ptni-3::HisCl1::SL2::tagBFP]* ; *krSi24[Punc-4::mNeonGreen]* | Multi-copy transgene inserted by irradiation |
| 3A | EN8086 | *krSi16[Pmyo-3::HisCl1::SL2::tagBFP]I* ; *egl-19(n582)IV* ; *krSi24[Punc-4::mNeonGreen]* |  |
| 3A | EN8087 | *krSi16[Pmyo-3::HisCl1::SL2::tagBFP]I* ; *unc-2(e55)X* ; *krSi24[Punc-4::mNeonGreen]* |  |
| 3A,B | EN8088 | *krSi16[Pmyo-3::HisCl1::SL2::tagBFP]I* ; *unc-36(e251)III* ; *krSi24[Punc-4::mNeonGreen]* |  |
| 3B | EN8423 | *krSi16[Pmyo-3::HisCl1::SL2::tagBFP]I* ; *unc-36(e251)III* ; *krSi24[Punc-4::mNeonGreen] ; krSi193[Punc-4::GFP-unc-36]* | miniMos single copy insertion |
| 3B | EN8424 | *krSi16[Pmyo-3::HisCl1::SL2::tagBFP]I* ; *unc-36(e251)III* ; *krSi24[Punc-4::mNeonGreen] ; krSi194[Pmyo-3::GFP-unc-36]* | miniMos single copy insertion |
| 3C | EN8873 | *krSi16[Pmyo-3::HisCl1::SL2::tagBFP]I* ; *[Punc-4::mNeonGreen] ; krIs89[Punc-4:: HisCat::SL2::tagBFP]* | Multicopy MosTI integration |
| 3C | EN8874 | *krSi16[Pmyo-3::HisCl1::SL2::tagBFP]I* ; *krSi24[Punc-4::mNeonGreen] ; krIs90[Punc-4:: HisCat::SL2::tagBFP]* | Multicopy MosTI integration |
| 3D | EN8787 | *krSi24[Punc-4::mNeonGreen] ; krSi257[Punc-4::HisCl1::SL2::tagBFP]I* ; | miniMos single copy insertion |
| 3D | EN8788 | *krSi16[Pmyo-3::HisCl1::SL2::tagBFP]I* ; *krSi24[Punc-4::mNeonGreen] ; krSi257[Punc-4::HisCl1::SL2::tagBFP]I* ; |  |
| 4A-C | EN3953 | *krSi16[Pmyo-3::HisCl1::SL2::tagBFP]I* ; *egl-3(gk238)V* ; *krSi24[Punc-4::mNeonGreen]* |  |
| 4A | EN7318 | *krSi16[Pmyo-3::HisCl1::SL2::tagBFP]I* ; *egl-3(gk238)V* ; *krSi24[Punc-4::mNeonGreen] ; krSi74 [Peft-3::egl-3(cDNA)::SL2::wScarlet]* | miniMos single copy insertion |
| 4B | EN7504 | *krSi16[Pmyo-3::HisCl1::SL2::tagBFP]I* ; *egl-3(gk238)V* ; *krSi24[Punc-4::mNeonGreen] ; krSi69 [Pmyo-3::egl-3(cDNA)::SL2::wScarlet]* | miniMos single copy insertion |
| 4C | EN7502 | *krSi16[Pmyo-3::HisCl1::SL2::tagBFP]I* ; *egl-3(gk238)V* ; *krSi24[Punc-4::mNeonGreen] ; krSi66 [Prgef-1::egl-3(cDNA)::SL2::wScarlet]* | miniMos single copy insertion |
| 6A | EN9504 | *krSi16[Pmyo-3::HisCl1::SL2::tagBFP]I* ; *flp-18(tm2179)X* ; *krSi24[Punc-4::mNeonGreen]* |  |
| 6B, E | EN9499 | *nlp-12(ok335)I krSi16[Pmyo-3::HisCl1::SL2::tagBFP]I* ; *krSi24[Punc-4::mNeonGreen]* |  |
| 6C | EN9501 | *krSi16[Pmyo-3::HisCl1::SL2::tagBFP]I* ; *nlp-11(yum480)II* ; *krSi24[Punc-4::mNeonGreen]* |  |
| 6D | EN9503 | *krSi16[Pmyo-3::HisCl1::SL2::tagBFP]I* ; *nlp-21(yum445)III* ; *krSi24[Punc-4::mNeonGreen]* |  |
| 6E | EN9644 | *nlp-12(ok335)I krSi16[Pmyo-3::HisCl1::SL2::tagBFP]I* ; *flp-18(tm2179)X* ; *krSi24[Punc-4::mNeonGreen]* |  |
| S1 | EN8216 | *krSi16[Pmyo-3::HisCl1::SL2::tagBFP]I* ; *egl-19(ad1006)IV* ; *krSi24[Punc-4::mNeonGreen]* |  |

Alleles in blue were generated by transgenesis; strains in black were obtained by genetic crosses. *Punc-4* is a short version (promC) of the *unc-4* promoter(Kratsios et al., 2015).
